# Histological transition during ovotestis formation in a female-to-male sex-change fish, the harlequin sandsmelt (*Parapercis pulchella*)

**DOI:** 10.1101/2025.08.25.672246

**Authors:** Akifumi Yao, Fumitaka Noguchi, Hisanori Kohtsuka, Toru Miura

## Abstract

Many teleost fishes are capable of sex change, and some species possess intersex gonads known as ovotestes, which contain both ovarian and testicular tissues. Uncovering the developmental processes in ovotestis formation is critical for elucidating the mechanisms of intersexuality in vertebrates. However, such developmental processes have been described in only a limited number of species. In this study, we examined the histological structure of juvenile gonads in the harlequin sandsmelt (*Parapercis pulchella*), a female-to-male sex-change fish in which mature females possess ovotestes. Juveniles measuring up to 32.2 mm in total length exhibited gonads containing cyst-formed developing oogonia and a small number of oocytes. In contrast, larger juveniles possessed more developed ovaries but showed no detectable spermatogenic germ cells. These results suggest that gonads initially differentiate as ovaries and subsequently develop into ovotestes through the emergence of male germ cells. Together with previous studies, our results also indicate that a similar developmental sequence may be shared among sex-change fishes.

## Introduction

Teleost fishes exhibit extraordinary diversity and plasticity in sexual patterns compared to other vertebrate lineages (Capel 2017; Nagahama et al. 2021). One particularly notable phenomenon is sex change, which has independently evolved in multiple teleost lineages (Kuwamura et al. 2020). Many sex-change fishes possess intersex gonads known as ovotestes, which contain both ovarian and testicular tissues but typically function as either ovaries or testes at a given time (Vega-Frutis et al. 2014; Yamaguchi and Iwasa 2017). Ovotestes are thought to facilitate rapid sex change by preparing tissues of the opposite sex in advance, which is beneficial for their reproduction (Kobayashi et al. 2005; Yamaguchi and Iwasa 2017).

In vertebrates, gonads generally differentiate precisely into either ovaries or testes through the antagonistic action of numerous feminizing and masculinizing factors (Capel 2017). Thus, uncovering the developmental processes of these gonads is essential for understanding how intersex traits can emerge in limited vertebrate species.

During early gonadal development in vertebrates, sexually undifferentiated gonads form first and subsequently differentiate into either ovaries or testes (Defalco and Capel 2009). Therefore, two possible developmental trajectories for ovotestis formation can be hypothesized: (1) undifferentiated gonads develop directly into ovotestes through the simultaneous initiation of oogenesis and spermatogenesis; or (2) undifferentiated gonads first differentiate into ovaries, and later transition into ovotestes with the onset of spermatogenesis.

In this study, we examined the harlequin sandsmelt, *Parapercis pulchella* (Temminck & Schlegel, 1843), a protogynous (female-to-male) sex-change fish, as a model to investigate the developmental mechanisms of ovotestis formation (Yao et al. 2023, 2024). While mature males form testes, mature females possess ovotestes (Hasebe 2019; Yao et al. 2023). However, the early stages of gonadal development remain unclear, due to difficulties in captive breeding and artificial fertilization. To address this knowledge gap, we conducted histological analyses of juvenile gonads using museum-preserved specimens to examine which of the two hypothesized developmental processes lead to ovotestis formation in the harlequin sandsmelt.

## Material and methods

Histological specimens of juvenile harlequin sandsmelt (10 individuals), preserved at the Tokai University Marine Science Museum (Shimizu, Shizuoka, Japan), were examined. Samples were selected based on the published catalog (Kobayashi 2004) and museum ledger. Analytical specimens were collected by SCUBA diving in Suruga Bay (off the coast of Kurumi, Numazu city), Shizuoka Prefecture, Japan on 16 October 1982. Then, the abdomens of these individuals were fixed using Bouin’s solutions, dehydrated with ethanol, clarified with xylene, embedded in paraffin, sectioned at 6–7 µm thickness, stained with hematoxylin and eosin, and preserved at that museum. Histological observations were conducted under an optical microscope equipped with a digital camera (Olympus BX51, DP74; Olympus, Tokyo, Japan). The sample information was obtained from the museum ledger and Kobayashi (2004) (summarized in **Table S1**). In addition, to confirm gonadal structures of adults in the same locality, we examined histological specimens from 19 adult gonads collected by hook and line in Suruga Bay (Enashi, Numazu City) in October 1979 (**Table S1**). Sex change stages, from female ovotestes to male testes, and developmental stages of germ cells were identified based on previous studies (Mandich et al. 2002; Qu et al. 2021; Yao et al. 2023).

## Results

Histological observations were conducted on juveniles (31.6–47.7 mm TL) and adults (97.9–163.8 mm TL) (**Fig. 1; Table S1**). Histological structures in sections were preserved despite over 40 years of storage, although hematoxylin staining was partially faded, especially at peripheral regions of sections (i.e., **Fig. 1e**).

**Fig. 1.**
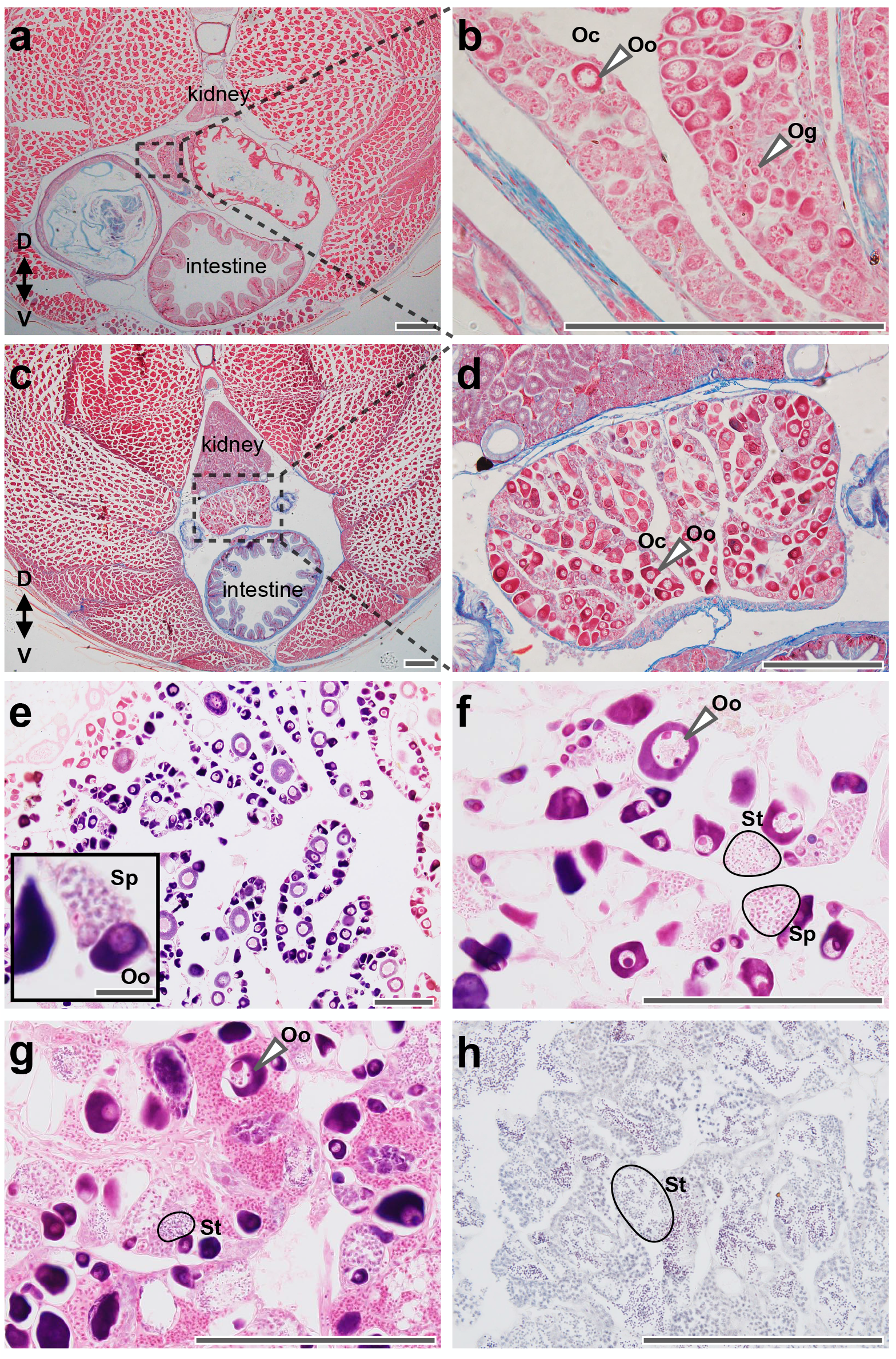
Gonad structure of juvenile harlequin sandsmelt (**a–d**) and mature individuals (**f-i**) from museum-preserved specimens. **a–b**. An individual of 31.6 mm TL. **c–d**. An individual of 47.7 mm TL. **a, c** Transverse section of the abdomen. **b, d** Higher-magnification view field shown in **a, c** respectively. Differentiated gonads contain oogonia and a few oocytes (perinucleolus stage). Immature ovaries contain numerous oocytes (perinucleolus stage) and oogonia. **e**. Female ovotestes. **f**. Early transition stage. **g**. Late transition stage. **h**. Male testes. Abbreviations: Og, oogonia; Oo, oocyte; Oc, ovarian cavity; Sp, spermatocyte; St, spermatid; D/V, dorsal-ventral axis. Scale bars = 200 and 20 µm (small panel in **e**).

In two relatively small juveniles (31.6–32.2 mm TL), developing ovaries consisted of many cysts-formed cell clusters with a few germ cells (around 10 µm diameter with a large nucleus). These cell cluster were considered to be oogonia. A small number of oocytes in the perinucleolus stage also appeared. Ovarian cavities had already been formed (**Fig. 1a–b**). Gonads in the remaining eight juveniles (32.8–47.7 mm TL) were larger than those of the small individuals with complexly branched ovarian cavities surrounded by ovarian lamellas. These gonads were identified as immature ovaries containing numerous perinucleolus-stage oocytes and oogonia. The number and size of oocytes were larger than those of the relatively small juveniles (**Fig. 1c–d**). Juveniles with testes were not found. Also, no histologically distinguishable male-type germ cells, such as spermatocytes, were observed in the gonads of any juveniles (**Fig. 1a–d**).

To confirm whether ovotestis formation in adult females and female-to-male sex change actually occur in this population, as previously reported in different populations (Hasebe 2019; Yao et al. 2023), we also examined the gonadal histology in adults collected at Suruga Bay. Gonads in all four defined sex change stages, including females that possessed ovotestes, sex-changing individuals, and male testes (defined by Yao et al. 2023), were found. Specifically, female ovotestes consisted of many oocytes along with a small number of spermatocyte clusters (**Fig. 1e**). Spermatids emerged in gonads undergoing sex change in the early transition stage (**Fig. 1f**). Testis tissues occupied more than half of the gonads and oocytes were degraded in the late transition stage (**Fig. 1g**). Male gonads were fully differentiated into testes, without any ovarian features (**Fig. 1h**).

## Discussion

This study revealed the early developmental process of gonads in the harlequin sandsmelt using museum-preserved histological specimens. Although gonads at the earliest developmental stages containing primordial germ cells were not observed, we identified juvenile gonads undergoing ovarian development (**Fig. 1a–d**). Furthermore, mature females collected from the same locality possessed ovotestes (**Fig. 1e–f**), consistent with previous studies from other localities (Hasebe 2019; Yao et al. 2023). These findings support the latter one of two proposed scenarios for ovotestis formation: (1) undifferentiated gonads directly develop into ovotestes; or (2) undifferentiated gonads first differentiate into ovaries and later transition into ovotestes. In addition, since no juveniles with testes were observed, this species is likely a monandric protogynous hermaphrodite, in which all males have changed sex from functional females (**Fig. 2**).

**Fig. 2.**
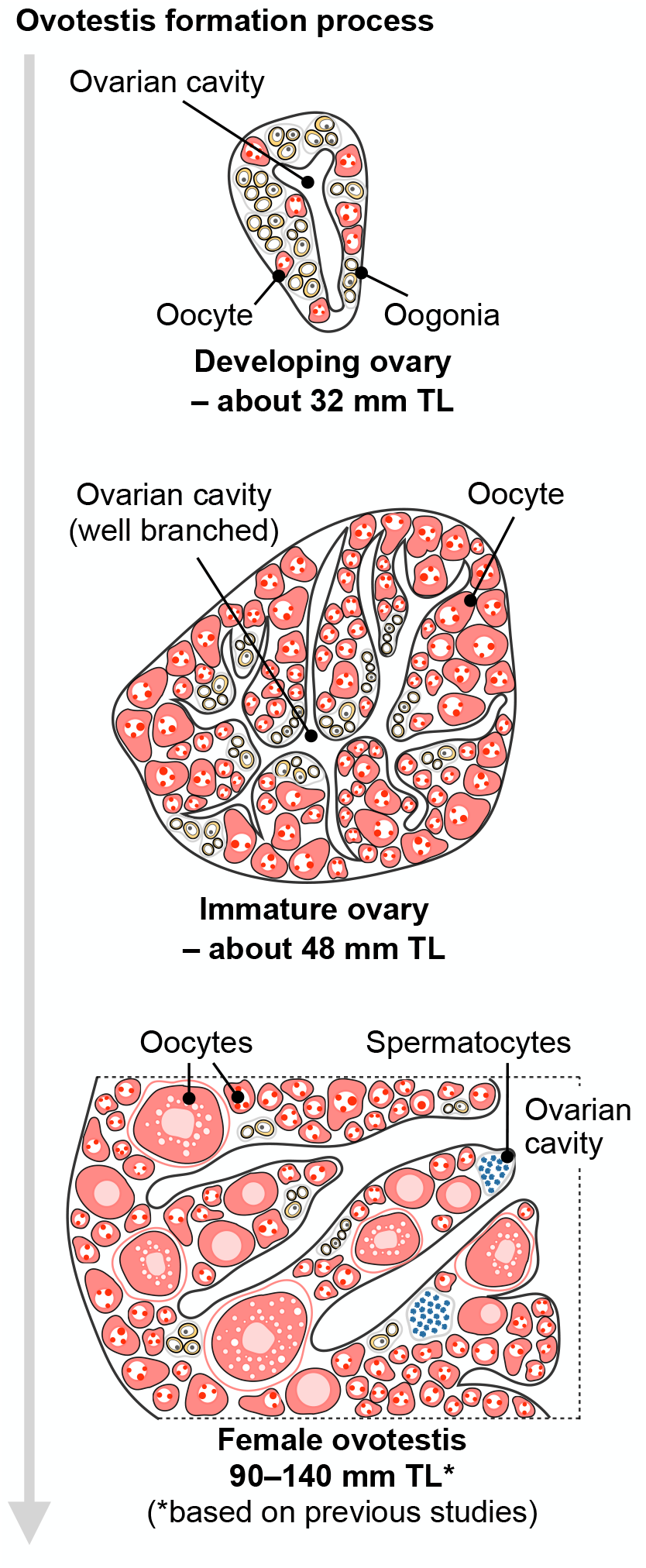
Schematic diagram of proposed ovotestis formation process in the harlequin sandsmelt. Gonads initially develop as ovaries and subsequently differentiate into female ovotestis through the onset of incomplete spermatogenesis.

Note that cysts composed of a few cells approximately 10 µm in diameter with large nuclei were observed in juveniles. These cells are presumed to be oogonia (Blazer 2002; Mandich et al. 2002). However, in teleosts, ovarian oogonia and testicular type A spermatogonia are morphologically similar and difficult to distinguish (Shang et al. 2018), and both exhibit high sexual plasticity (Lacerda et al. 2014). Therefore, further characterization using molecular markers will be necessary to clarify their identity.

This study utilized histological sections prepared over 40 years ago and deposited in the museum. Although some samples showed degradation, many were sufficiently preserved for observation. The museum houses an extensive collection of histological specimens of fish gonads, with detailed metadata (Kobayashi 2004). Thus, this resource is useful for research in fish physiology, developmental biology, and aquaculture.

Together with previous studies, our findings suggest that similar gonadal developmental processes may be shared across sex-change fishes from distinct lineages. For instance, in the U-mark sandperch, *Parapercis snyderi*, a comparable ovotestis development pattern has been reported (Kobayashi et al. 1993). The genus *Parapercis* includes protogynous species both with and without ovotestes in females, as well as gonochoristic species (Kobayashi et al. 1993; Randall 2003; Yao et al. 2023). Therefore, the evolutionary processes of ovotestis in the genus *Parapercis* should be carefully evaluated in further studies.

Similar developmental trajectories have been observed in species belonging to other orders. In the yellowspotted rockcod, *Epinephelus areolatus*, mature females possess ovotestes, while smaller, immature females form ovaries (Boddington et al. 2021), suggesting parallel evolution of ovotestis formation in different protogynous species. Moreover, in bidirectional sex-change species such as the yellow hawkfish, *Cirrhitichthys aureus*, and in protandrous (male-to-female sex change) species (e.g., the yellowtail clownfish, *Amphiprion clarkii*, and the black porgy, *Acanthopagrus schlegelii*), gonads initially differentiate into ovaries, which later transition into either female or male ovotestes (Kobayashi and Suzuki 1992; Lee et al. 2001; Miura et al. 2003). These findings suggest that ovotestis formation may follow a shared developmental process across sex-change fishes, regardless of phylogenetic position or the direction of sex change. It can be considered that shared developmental processes might reflect underlying developmental constraints.

In conclusion, this study reveals the histological features of juvenile gonads in the harlequin sandsmelt and uncovers the developmental process of ovotestis formation in this species, providing an experimental basis for elucidating molecular mechanisms of ovotestis formation. We also highlight a shared ovotestes formation process across sex-change fishes. These findings provide insights into the developmental mechanisms and evolutionary processes of ovotestes in broader teleost lineages, leading to a better understanding of intersexuality in vertebrates.

## Acknowledgments

We express our gratitude to the late Dr. Koji Kobayashi (Tokai University Marine Science Museum) for his valuable contributions to the preparation and maintenance of specimens used in this study for a long period. We thank Mr. Yoshiaki Uchida (The University of Tokyo) and the staff of Tokai University Marine Science Museum for their kind support for observations of specimens.

## Author contributions

AY and TM designed this study. AY, FN and HK carried out observations. AY and TM analyzed data and wrote the first draft of this manuscript. All authors read and approved the final manuscript.

## Notes

### Competing Interest Statement

The authors have declared no competing interest.

## References

Blazer VS (2002) Histopathological assessment of gonadal tissue in wild fishes. Fish Physiol Biochem 26:85–101. 10.1023/A:1023332216713

Boddington DK, Wakefield CB, Fisher EA, et al (2021) Age, growth and reproductive life-history characteristics infer a high population productivity for the sustainably fished protogynous hermaphroditic yellowspotted rockcod (Epinephelus areolatus) in north-western Australia. J Fish Biol 99:1869–1886. 10.1111/jfb.14889

Capel B (2017) Vertebrate sex determination: Evolutionary plasticity of a fundamental switch. Nat Rev Genet 18:675–689. 10.1038/nrg.2017.60

Defalco T, Capel B (2009) Gonad Morphogenesis in Vertebrates: Divergent Means to a Convergent End. Annu Rev Cell Dev Biol 25:457–482. 10.1146/annurev.cellbio.042308.13350

Hasebe K (2019) Reproductive ecology and sex change in two Pinguipetidae species at Tateyama Bay. Tokyo University of Marine Science and Technology

Kobayashi K (2004) Catalogue of the collections deposited in marine science museum, Tokai University. Sci Reports Museum, Tokai Univ 6:35–111

Kobayashi K, Suzuki K (1992) Hermaphroditism and sexual function in Cirrhitichthys aureus and the other japanese hawkfishes (Cirrhitidae: Teleostei). Japanese J Ichthyol 38:397–410

Kobayashi K, Suzuki K, Shiobara Y (1993) Reproduction and hermaphroditism in Parapercis snyderi (Teleostei, Parapercidae) in Suruga Bay, central Japan. J Fac Mar Sci Technol Tokai Univ 35:149– 168

Kobayashi Y, Sunobe T, Kobayashi T, et al (2005) Gonadal structure of the serial-sex changing gobiid fish Trimma okinawae. Dev Growth Differ 47:7–13. 10.1111/j.1440-169x.2004.00774.x

Kuwamura T, Sunobe T, Sakai Y, et al (2020) Hermaphroditism in fishes: an annotated list of species, phylogeny, and mating system. Ichthyol Res 67:341–360. 10.1007/s10228-020-00754-6

Lacerda SMN, Costa GMJ, França LRDe (2014) Biology and identity of fish spermatogonial stem cell. Gen Comp Endocrinol 207:56–65. 10.1016/j.ygcen.2014.06.018

Lee YH, D. JL, Yueh WS, et al (2001) Sex change in the protandrous black porgy, Acanthopagrus schlegeli: A review in gonadal development, estradiol, estrogen receptor, aromatase activity and gonadotropin. J Exp Zool 290:715–726. 10.1002/jez.1122

Mandich A, Massari A, Bottero S, Marino G (2002) Histological and histochemical study of female germ cell development in the dusky grouper Epinephelus marginatus (Lowe, 1834). Eur J Histochem 46:87– 100. 10.4081/1657

Miura S, Komatsu T, Higa M, et al (2003) Gonadal sex differentiation in protandrous anemone fish, Amphiprion clarkii. Fish Physiol Biochem 28:165–166. 10.1023/B:FISH.0000030513.05061.88

Nagahama Y, Chakraborty T, Paul-Prasanth B, et al (2021) Sex determination, gonadal sex differentiation, and plasticity in vertebrate species. Physiol Rev 101:1237–1308. 10.1152/physrev.00044.2019

Qu M, Cao X, Wang H, et al (2021) Gonadal structure and expression localization of sex-related genes in the hermaphroditic grouper Epinephelus akaara (Perciformes: Epinephelidae). Aquaculture 542:736902. 10.1016/j.aquaculture.2021.736902

Randall JE (2003) Review of the sandperches of the Parapercis cylindarica complex (perciformes: pinguipetidae) with description of two new species from the western pacific. Bish museum Occas Pap 72:1–19

Shang M, Su B, Perera DA, et al (2018) Testicular germ line cell identification, isolation, and transplantation in two North American catfish species. Fish Physiol Biochem 44:717–733. 10.1007/s10695-018-0467-3

Vega-Frutis R, Macías-Ordóñez R, Guevara R, Fromhage L (2014) Sex change in plants and animals: A unified perspective. J Evol Biol 27:667–675. 10.1111/jeb.12333

Yamaguchi S, Iwasa Y (2017) Advantage for the sex changer who retains the gonad of the nonfunctional sex. Behav Ecol Sociobiol 71:39. 10.1007/s00265-017-2269-5

Yao A, Kohtsuka H, Miura T (2024) Reference transcriptome assembly of a protogynous sex change fish, harlequin sandsmelt (Parapercis pulchella). Mar Genomics 73:101086. 10.1016/j.margen.2024.101086

Yao A, Nakamura M, Kohtsuka H, et al (2023) Gonadal and cellular dynamics during protogynous sex change in the harlequin sandsmelt Parapercis pulchella. J Fish Biol 103:1347–1356. 10.1111/jfb.15534

