## Supplementary material for "Histological transition during ovotestis formation in a female-to-male sex-change fish, the harlequin sandsmelt (*Parapercis pulchella*)": Table S1 (caption)

**Caption of supplemental table**

**Online Resource 2 (Table S1).**

**Data for analyzed specimens preserved at the Tokai Marine Science Museum.**

These specimen’s metadata were obtained from the museum ledger and Kobayashi (2004). This table included 15 juveniles and 20 adults (five juveniles and one adult were observed, but not included in this study due to poor quality of histological specimens). Sample ID were described both in the museum ledger and Kobayashi (2004). Lot number was assigned in Kobayashi (2004). Specimens could be identified using these numbers. Sampling sites and sampling methods were noted in Kobayashi (2004). Total length and standard length of each specimen were recorded in the museum ledger. Histological features were described based on the present study.

### Citations

Kobayashi K (2004) Catalogue of the collections deposited in marine science museum, Tokai University. Sci Reports Museum, Tokai Univ 6:35–111 https://www.muse-tokai.jp/wp/wp-content/uploads/2017/09/bulletin_06.pdf
